# Visual statistical learning alters low-dimensional cortical architecture

**DOI:** 10.1101/2023.10.24.563271

**Authors:** Keanna Rowchan, Daniel J. Gale, Qasem Nick, Jason P. Gallivan, Jeffrey D. Wammes

## Abstract

Our brains are in a near constant state of generating predictions, extracting regularities from seemingly random sensory inputs to support later cognition and behavior – a process called statistical learning (SL). Yet, the activity patterns across cortex and subcortex that support this form of associative learning remain unresolved. Here we use human fMRI and a visual SL task to investigate changes in neural activity patterns as participants implicitly learn visual associations from a sequence. By projecting functional connectivity patterns onto a low-dimensional manifold, we reveal that learning is selectively supported by changes along a single neural dimension spanning visual-parietal and perirhinal cortex (PRC). During learning, visual cortex expanded along this dimension, segregating from other networks, while dorsal attention network (DAN) regions contracted, integrating with higher-order transmodal cortex. When we later violated the learned associations, PRC and entorhinal cortex, which initially showed no evidence of learning-related effects, now contracted along this dimension, integrating with the default mode and DAN, while decreasing covariance with visual cortex. Whereas previous studies have linked SL to either broad cortical or medial temporal lobe changes, our findings suggest an integrative view, whereby cortical regions reorganize during association formation, while medial temporal lobe regions respond to their violation.

## INTRODUCTION

Each day, we are inundated with an influx of sensory information. Despite the apparent randomness of these inputs, we possess the remarkable ability to discern repeating patterns in our environment. Our ability to implicitly extract regularities over time and across development^1^ is known as statistical learning (SL), and it enables us to make predictions about the world to inform our decisions and behavior^2,3^. One well studied form of such learning is visual SL, which examines our ability to internally represent and extract probabilities for temporally co-occurring visual stimuli^4,5^.

Traditionally, SL was thought to engage primarily lower-order task-specific sensory cortices, whereas the medial temporal lobe (MTL; including hippocampus, entorhinal cortex (ERC), perirhinal cortex (PRC), and parahippocampal cortex) was mainly associated with episodic learning and memory^6,7^. In recent years, however, these neural distinctions have eroded in light of new evidence. Specifically, it has been shown that, in addition to the sensory-specific cortices, SL recruits higher-order association networks^8,9^, suggesting that it requires both task-specific ^10–12^ and task-general cognitive processing^13–15^. Recent studies have also shown that the MTL and hippocampus exhibit representational change and differential responses during SL^16–18^, and that hippocampal volume predicts SL performance^19,20^. Further challenging the traditional view, patient work has shown that patients with damage to the hippocampus and surrounding MTL show little evidence for SL,^21^ suggesting a critical role for these regions.

While these newer results align with the MTL’s established role in learning^22,23^ and its sensitivity to temporal and spatial information^24,25^, they nevertheless indicate that the traditional distinctions between cortical and MTL processes require updating. One advance in this area has been to propose a division of labor within the hippocampal system, where the monosynaptic pathway (between the ERC and CA1) supports SL, and the trisynaptic pathway (between the ERC, CA1, CA3, and dentate gyrus, DG) supports episodic learning^26^. Consistent with these complementary roles, direct stimulation in the monosynaptic pathway (CA1) impairs SL^27^, but focal trisynaptic pathway (DG) lesions do not^28^. Although this new evidence highlights the role of the MTL in SL, there remains some uncertainty. While the patient work described above had found MTL lesions impaired SL, some patients with extensive bilateral hippocampal lesions show evidence of, and at times, outperform controls in SL^15,29,30^. These conflicting findings leave the relative contributions of the cortex and MTL to SL unresolved.

While many individual brain areas have been identified as contributors to SL^8,12,15^, it is clear that such learning involves the coordination and integration of information processing across multiple distributed brain regions^31–33^. Indeed, while prior work highlights the MTL, task-specific, and task-general cortical networks in SL, each of these areas are themselves reciprocally interconnected across several distributed, whole-brain networks^34^. Understanding these interconnections requires the study of functional connectivity between brain areas embedded within large-scale networks, and analytical approaches capable of effectively characterizing the changes in whole-brain functional architecture that occur during SL. Such an approach is crucial not only for characterizing whole-brain interactions but also for asking critical questions about the relative extent and timing of connectivity changes during SL and how these systems independently or cooperatively encode regularities^16,29,30^.

Here we adopt a novel approach for effectively characterizing large-scale patterns of whole-brain functional activity along a low-dimensional subspace or manifold, which reflects the main patterns of covariance across brain regions^35–37^. In other work, this approach has not only provided fundamental insights into the principles governing the activity of large-scale neural populations in many brain areas^38,39^, but also the intrinsic functional organization of the brain in both healthy and patient populations^40,41^. Here, we adopted these methods to understand how cortical and MTL areas functionally interact at various stages of visual SL, providing a window into the evolving landscape of brain activity during SL and how these different brain areas adapt to changing environmental patterns.

## RESULTS

The current study analyzed previously published functional magnetic resonance imaging (fMRI) data^18^ from participants (N = 33) performing a visual SL task, where they viewed abstract images presented one at a time (see Fig. 1A). The task included eight blocks, where participants viewed a sequence of 80 images consisting of 5 repetitions of 16 unique images, totalling 640 image presentations. During six of these blocks (blocks 2-7), the sequences contained embedded regularities, such that one image (A) was always followed by its pairmate (B; See Fig. 1A, 1B left). These SL blocks were bookended by two, unstructured blocks (blocks 1 and 8), where images were presented in a random order (see Fig. 1B right). Importantly, participants were unaware of this covertly embedded structure, and engaged in a basic perceptual task to maintain their attention, which was completely orthogonal to the pair structure^18^ (see Materials and Methods).

**Figure 1.**
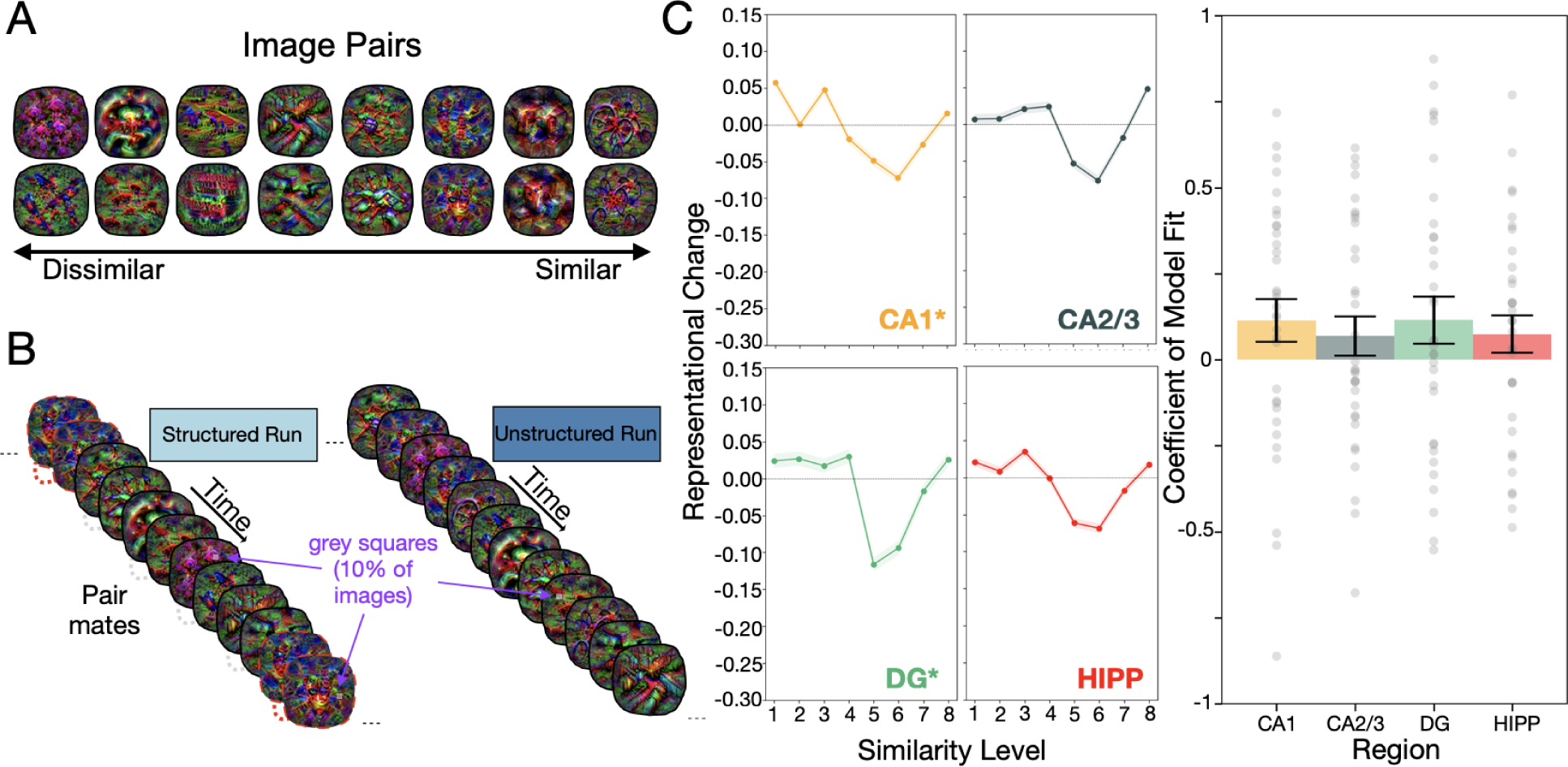
Overview of visual SL task and representational change predicted by the nonmonotonic plasticity hypothesis. (A) Pairmates used in the visual SL task^18^, ordered from most visually dissimilar, to most visually similar. (B) Visual SL task structure. Participants viewed images presented one at a time on screen. Structured runs contained an embedded pair structure, whereby an A image was always followed by its pairmate (B image; example pairmates denoted by red dashed line). By contrast, unstructured runs presented images in a random order. Each run consisted of 80 image presentations (5 repetitions of each pair). Participants were tasked with pressing a button when a small faded gray square appeared in the image (10% of trials). (C) Left, Relationship between image similarity and representational change predicted by the nonmonotonic plasticity hypothesis. For each region (CA1, CA2/3, DG and whole hippocampus; HIPP), we correlated parameter estimates derived from a general linear model for each A and B image in a pair, and created comparison scores by subtracting Pre-from Post-learning correlation scores. This measure quantifies the degree of representational change between image pairs as participants underwent visual SL (i.e., from Early-to Late-learning), with positive scores indicating integration, and negative scores indicating differentiation. The comparison scores were plotted as a function of similarity level (i.e., coactivation) for each of the image pairs (ordered from least to greatest visual similarity in pairmates). Shaded areas depict bootstrap resampled standard errors of the means. Right, theory-constrained cubic models were fit to the representational change patterns in all but one held-out participant, then used to predict the held-out participants’ scores. These individual-specific model fits are shown in the light gray scatter plots. Barplots display the group-level model fit, measured by the correlation between actual and predicted values. At a group-level, representational change was reliably predicted by a theory-constrained U-shaped model in the CA1 and DG. Error bars depict SEM.

### Statistical learning results in neural changes at the regional level

Before examining whole-brain changes in functional connectivity, we first verified that subjects exhibited learning during the SL blocks. Prior work, including our own, has shown that visual SL leads to changes in stimulus-specific patterns of neural activity in the hippocampus^16–18^. Depending on the visual similarity between pairmates, learning can either lead to neural integration, where two temporally coupled stimuli become more similar in their neural representations, or to neural differentiation, whereby they become less similar. Recent theoretical^42^, modeling^43^, and empirical^18,44^ work indicates this relationship follows a U-shaped function, as described by the nonmonotonic plasticity hypothesis^42^ (NMPH). In our prior work^18^ the strength of this U-shaped function in the DG provided the primary measure that SL had taken place. Consistent with these findings, our re-analysis of the current data subset used for our connectivity analyses (see Supplementary Extended Materials and Methods) revealed a U-shaped learning effect in the CA1 (r=0.11, 95% CI=[-0.010, 0.232], p=0.0351) and DG (r=0.11, 95% CI=[-0.017, 0.248], p=0.0451), but not the CA2/3 (r=0.07, 95% CI=[-0.044, 0.180], p=0.115), or across the entire hippocampus when subfields were combined (r=0.07, 95% CI=[-0.032, 0.180], p=0.088; see Fig. 1C). These results replicate prior effects^18^ demonstrating SL-induced representational changes in the hippocampus. With this demonstration of learning now in place, we next examined whether, and to what extent, functional interactions both within and outside of the MTL system are altered during SL.

### Investigating changes in functional brain organization during statistical learning

To investigate changes in whole-brain functional organization during visual SL, we selected four functional runs, each representing a key epoch of the task: Early- and Late-learning epochs (the first and last structured blocks, runs 2 and 7), and Pre- and Post-learning epochs (the first and last unstructured blocks, runs 1 and 8). For each participant and epoch, we extracted the mean blood oxygenation level-dependent (BOLD) timeseries data for each cortical region pre-defined by the Schaefer-1000 parcellation^45^, and MTL regions using individual-specific parcellations generated using the Automated Segmentation of Hippocampal Subfields software package^46^ (ASHS). We then computed covariance matrices (representing functional connectivity between each pairwise set of regions) for each of our task epochs (Pre-, Early-, Late-, and Post-learning). [see Fig. 2A; see Supplementary Extended Materials and Methods].

**Figure 2.**
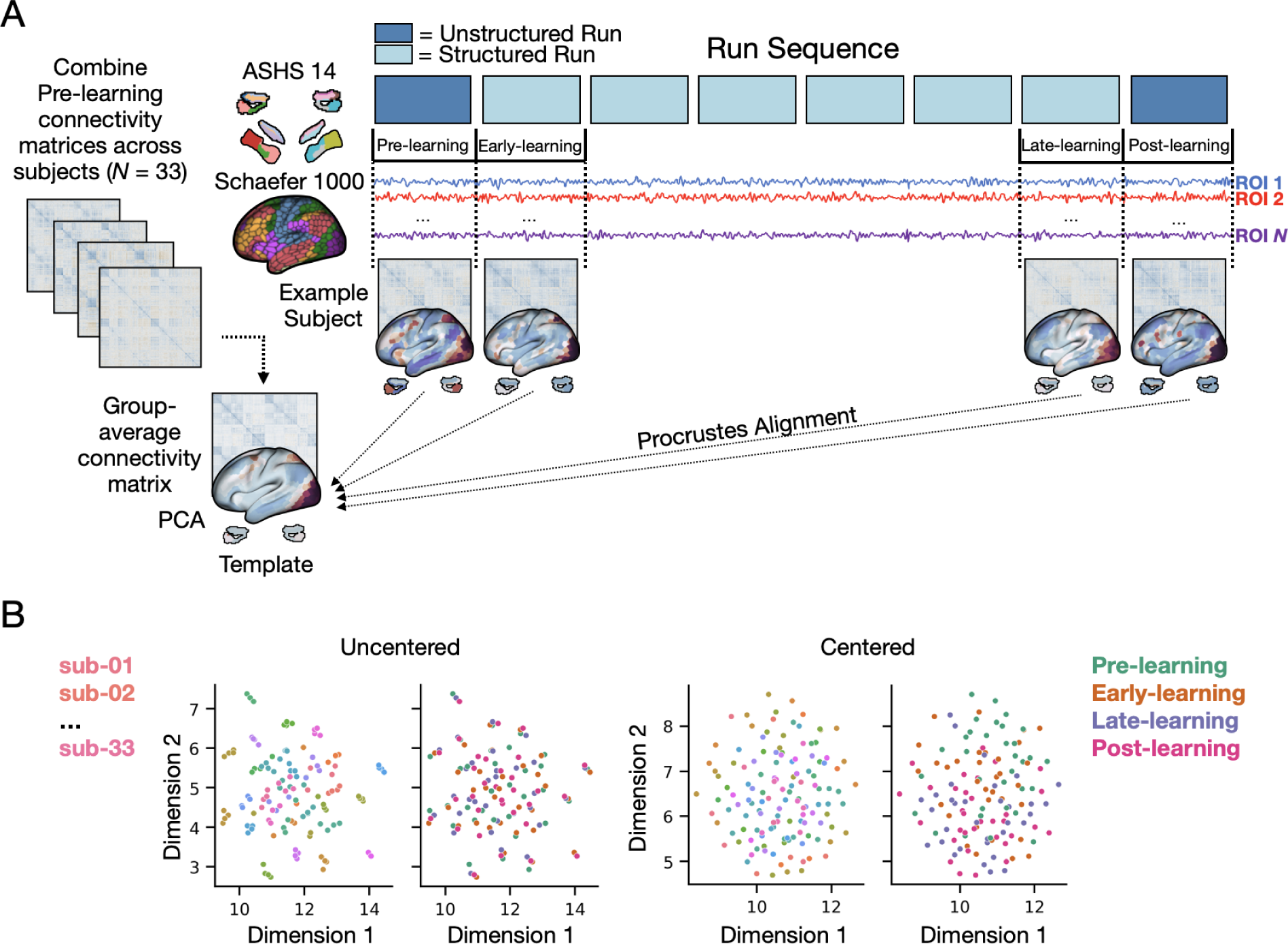
Overview of neural analytic approach. (A) Functional connectivity analysis approach. For each participant and epoch (Pre-, Early-, Late-, and Post-learning), we computed functional connectivity matrices from timeseries data extracted from the Schaefer-1000 cortical parcellation^45^ and participant-specific hippocampal parcellations derived from the Automated Segmentation of Hippocampal Subfields^46^ software. We then estimated functional connectivity manifolds for each epoch using PCA with centered and thresholded connectivity matrices (see Materials and Methods). All manifolds were aligned to a common Pre-learning Template manifold using Procrustes alignment. (B) Uniform Manifold Approximation^49^ visualization of connectivity matrices, both before and after centering. Single data points represent individual matrices. For both uncentered and centered data, the left plots have the data points colored according to subject identity whereas the right plots have the data points colored according to task epoch. Prior to centering, matrices were mainly clustered according to subject identity whereas this cluster structure was abolished following centering.

Prior work indicates that individual differences account for the majority of variance in patterns of brain activity, often obscuring task-related changes in functional connectivity^35,47^. We recently developed a Riemannian manifold centering approach^47^ to mitigate these effects, making task-related changes in brain structure more readily detectable^48^ (see Methods and Materials). To illustrate the impact of this method, we projected participants’ covariance matrices both before and after this centering procedure using uniform manifold approximation^49^ (UMAP). Consistent with prior work, including our own^35,50,51^, before centering, we found that the covariance matrices mainly clustered according to subject identity (see Fig. 2B left). However, after centering, this subject-level clustering was completely abolished, thus permitting exploration of task-related changes in connectivity across epochs (see Fig. 2B right).

Next, we used these centered matrices to estimate cortical-subcortical connectivity manifolds for each participant’s Pre-, Early-, Late-, and Post-learning epochs. Building from prior work^35,50,52^, we converted each matrix into an affinity matrix before applying principal components analysis (PCA) to reveal sets of principal components (PCs) that provide a low-dimensional representation of cortical-MTL structure. Finally, we aligned each manifold to a common template manifold, constructed using the mean of all participants’ Pre-learning matrices (see Fig. 2A). This alignment served two key purposes: First, it provided a common target for manifold alignment, such that all manifolds could be compared within a common neural space, and second, it allowed us to directly examine learning-specific deviations away from this Pre-learning template manifold, thus enhancing our sensitivity to detect learning-related changes in manifold structure^35^.

### Cortical and subcortical manifold structure prior to learning

The top three PCs of our Pre-learning template manifold describe the main dimensions of functional connectivity observed during the task (see Fig. 3A), and explain the majority of the variance in the data (60.0%; see Fig 3B). PC1 distinguishes the visual network, parietal dorsal attention network (DAN), and PRC (regions in red) from other cortical regions (in blue). PC2 separates transmodal, default mode network (DMN) regions (in red) from somatomotor regions (in blue). Lastly, PC3 distinguishes medial visual cortex (in red) from areas of the DAN (in blue). Together, PC1 and PC2 differentiate visual and somatomotor regions from transmodal regions of the DMN (see Fig. 3C, left and middle), reflecting the brain’s intrinsic functional architecture^36,37^. This tripartite architecture is thought to reflect a crucial feature of cortical brain organization, where the shift from unimodal to transmodal networks signifies a global processing hierarchy ascending from lower-order sensory and motor systems, to higher-order association areas of the cortex, such as the DMN^53^. To visualize this tripartite structure, we projected each brain region’s loadings on all three PCs onto a three-dimensional manifold space (see Fig. 3D).

**Figure 3.**
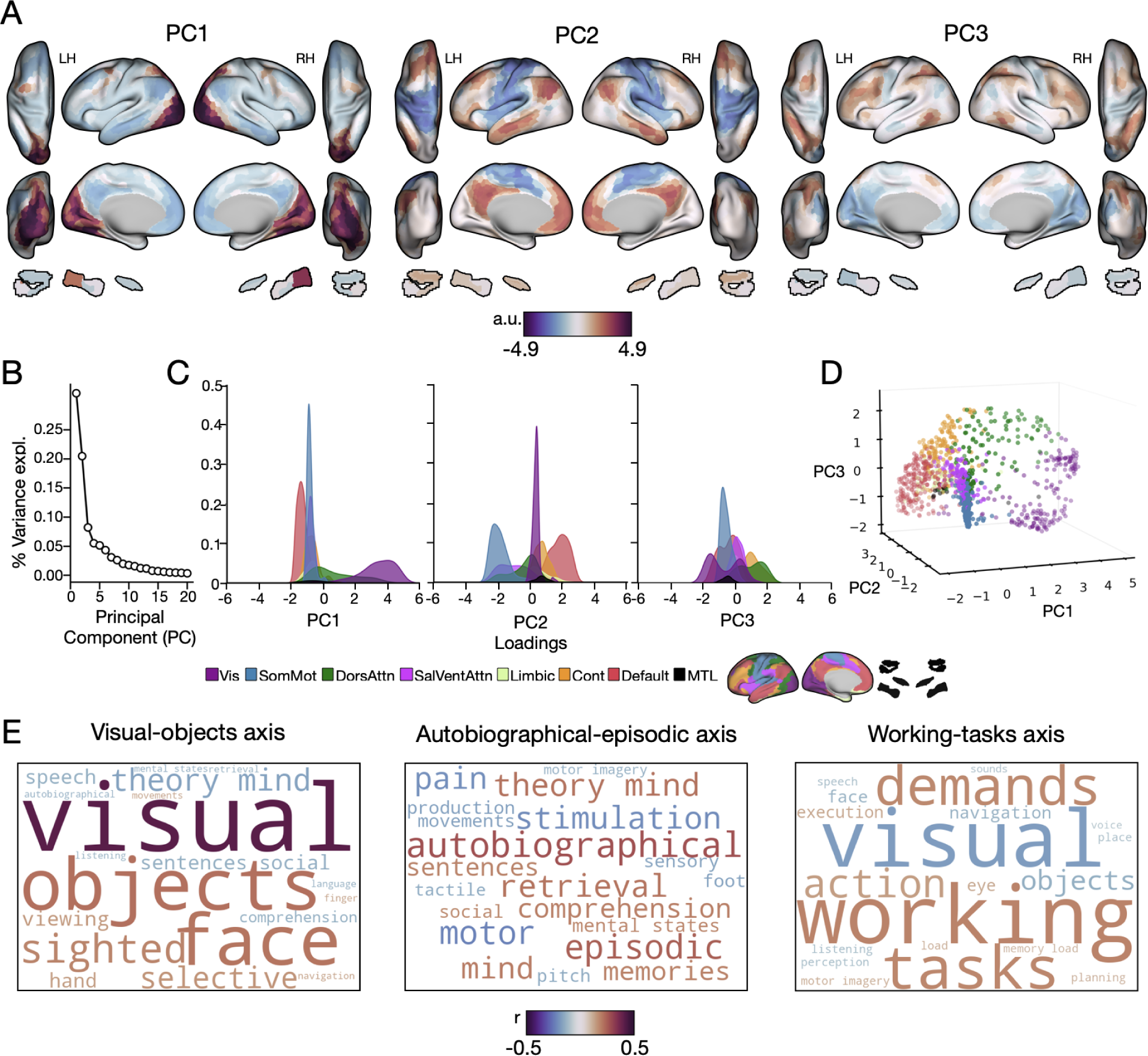
Functional organization of the Pre-learning epoch. (A) Cortical-Subcortical loadings for each of the top three principal components (PCs), projected in surface space. (B) Percent variance explained for the first 20 PCs. (C) Histograms displaying the distributions of each network’s loading along each PC, colored according to its functional network assignment, denoted on the bottom^55^. (D) Functional organization of the Pre-learning epoch. Each brain region is depicted as a point in a three-dimensional manifold space, with its location determined by its loading on the first three PCs. (E) Meta-analysis of PC loadings submitted to the Neurosynth Decoder tool^54^. Word clouds show the top and bottom ten functions likely associated with each PC. The text size and colour within each word cloud describes their correlation value (bigger words have higher correlation values and smaller words have smaller values) and their polarity (words in red are more positive and words in blue are more negative).

To better understand the typical functions associated with each of our PCs, we conducted a meta-analysis using the Neurosynth Decoder^54^. This tool quantifies patterns of whole-brain activity against a comprehensive dataset of whole-brain maps from thousands of fMRI studies. The results of this analysis are in the form of correlation values, revealing associations between each PC map, and various keywords which often reflect tasks or cognitive functions. For the purposes of visualization, we transformed these correlation values into word clouds (see Fig. 3E), where the color and size of a keyword represents the direction and magnitude of the correlation. We used these word clouds to assign meaningful functional labels to each PC. We termed PC1 the visual-objects axis, emphasizing the role of its constituent regions in object recognition and visual processing; PC2 the autobiographical-episodic axis, given the role of its constituent regions in memory and time-related cognition; and PC3 the working-tasks axis, given the role of its constituent regions in multiple tasks.

While together, the top three PCs recreate the tripartite structure seen in prior work^35–37^ (see Fig. 3D), the order of variance explained by these orthogonal dimensions offers insights into the specific neural demands of our task. Notably, prior resting-state studies using the same dimension reduction method (PCA) have identified our visual-object connectivity axis (PC1) as explaining the third-highest proportion of variance, while PC2 and PC3 commonly explain the first and second highest variances, respectively^35,36^. However, considering that the stimuli in our task were originally designed to engage higher-order object-selective visual regions^18^, it is intuitive that PC1 explains the greatest amount of variance in our data. Given this, we next examined whether visual SL selectively modulated connectivity motifs along this visual-object axis.

### Changes along the visual-object connectivity axis during statistical learning

Figure 4A shows the average loading of each brain region along the visual-object (PC1) and autobiographical-episodic (PC2) axes as a function of task epoch. Notably, Late-learning appeared to be associated with a compression along the visual-object axis, which then expanded along its positive loading direction once SL was interrupted during Post-learning. To test whether this pattern of compression and expansion along the visual-objects axis was significant across task epochs, we calculated, for each participant, the min-max spread along this axis for each of our four task epochs (Pre-, Early-, Late-, and Post-learning) and performed a repeated measures ANOVA (rmANOVA) on these values. This analysis revealed a significant effect of epoch along the visual-objects axis (*F*(3, 96)=5.10, p=0.008), indicating changes (contraction and expansion) in the distribution of regional embeddings along this axis during SL (for a similar approach see^40^). Notably, when we performed this same analysis for the autobiographical-episodic (PC2) and working-task (PC3) axes, we observed no significant changes in expression across epochs (autobiographical-episodic axis: *F*(3, 96)=0.61, p=0.61; working-task axis: *F*(3, 96)=2.47, p=0.10), suggesting a task-related selectivity to these effects.

Fig. 4B highlights pairs of brain regions (left and right hemisphere) from three distinct networks that load positively onto the visual-objects axis (visual network: purple; DAN: green; PRC: black). Together, these regions indicate that there appear to be large, yet differentiable shifts in the embedding of certain regions across epochs. For example, the visual network and DAN exhibit large shifts from Early- to Late-learning (see Fig. 4B middle), whereas the PRC exhibits its largest shifts from Late- to Post-learning (see Fig. 4B right). To test for these and any other regional effects, we performed a rmANOVA for each region’s loadings onto the visual-object axis across task epochs. Following FDR-correction^56^ (q < 0.05), we found that 129 regions exhibited a significant main effect of epoch (see Fig. 5A). These regions were within two major cortical networks: the visual network (63 regions) and DAN (37 regions); with smaller clusters also found in the DMN (9 regions), cingulo-opercular network (8 regions), somatomotor network (6 regions), limbic network (2 regions), and Salience and Ventral Attention (SalVentAttn) network (1 region). Notably, we also found significant areas in the MTL, including left and right PRC, and right ERC (see Fig. 5A). Figure 5B plots the trajectory of change for significant regions along the visual-objects axis.

**Figure 4.**
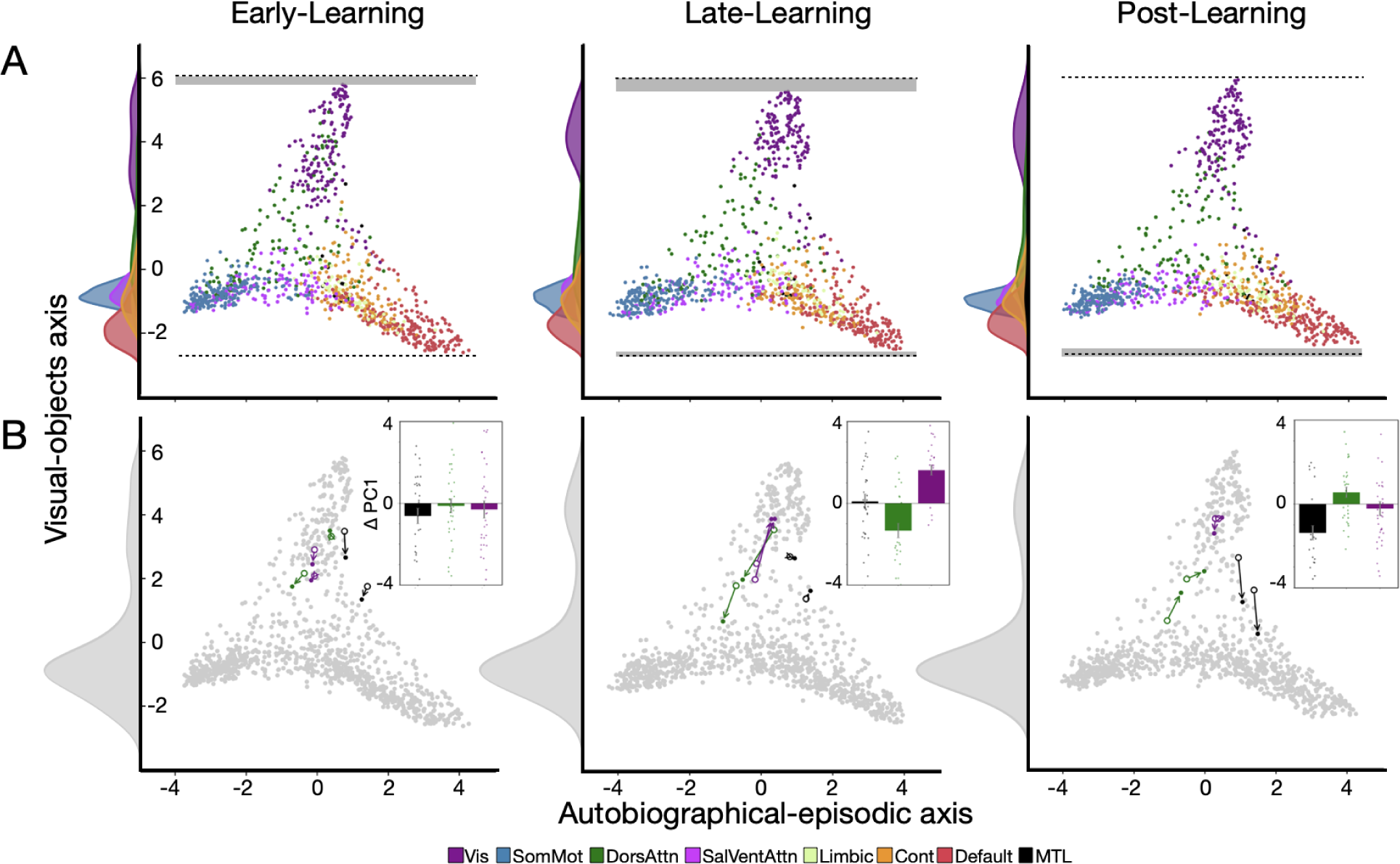
Changes in group-average manifold loadings along the visual-objects axis. (A) Changes in functional organization during Early-, Late- and Post-learning epochs. Each brain region is shown as a point in a two-dimensional manifold space along the visual-objects (PC1) and autobiographical-episodic (PC2) axis, with its color determined by its functional network assignment, denoted at the bottom^55^. The horizontal dashed line and bars across epochs (in grey) allows for a direct visual comparison of the change in the expression of the visual-objects axis across task epochs. Histograms along the Y-axis show point densities for each epoch and functional network. (B) Temporal trajectories represent left and right hemisphere seed regions. The starting position of each seed region from the preceding epoch is marked by an open white circle, with arrows indicating the displacement of each region in the current epoch (colored-in endpoint^55^). Gray histograms along the Y-axis display the mean density of points across all regions. Inset bar plots within each task epoch display the average change (Δ) in loading for each pair of seed regions from the preceding to current epoch. Negative and positive values reflect decreases and increases along the visual-objects axis, respectively. Single data points depict single subjects. Error bars depict standard error of the mean.

**Figure 5.**
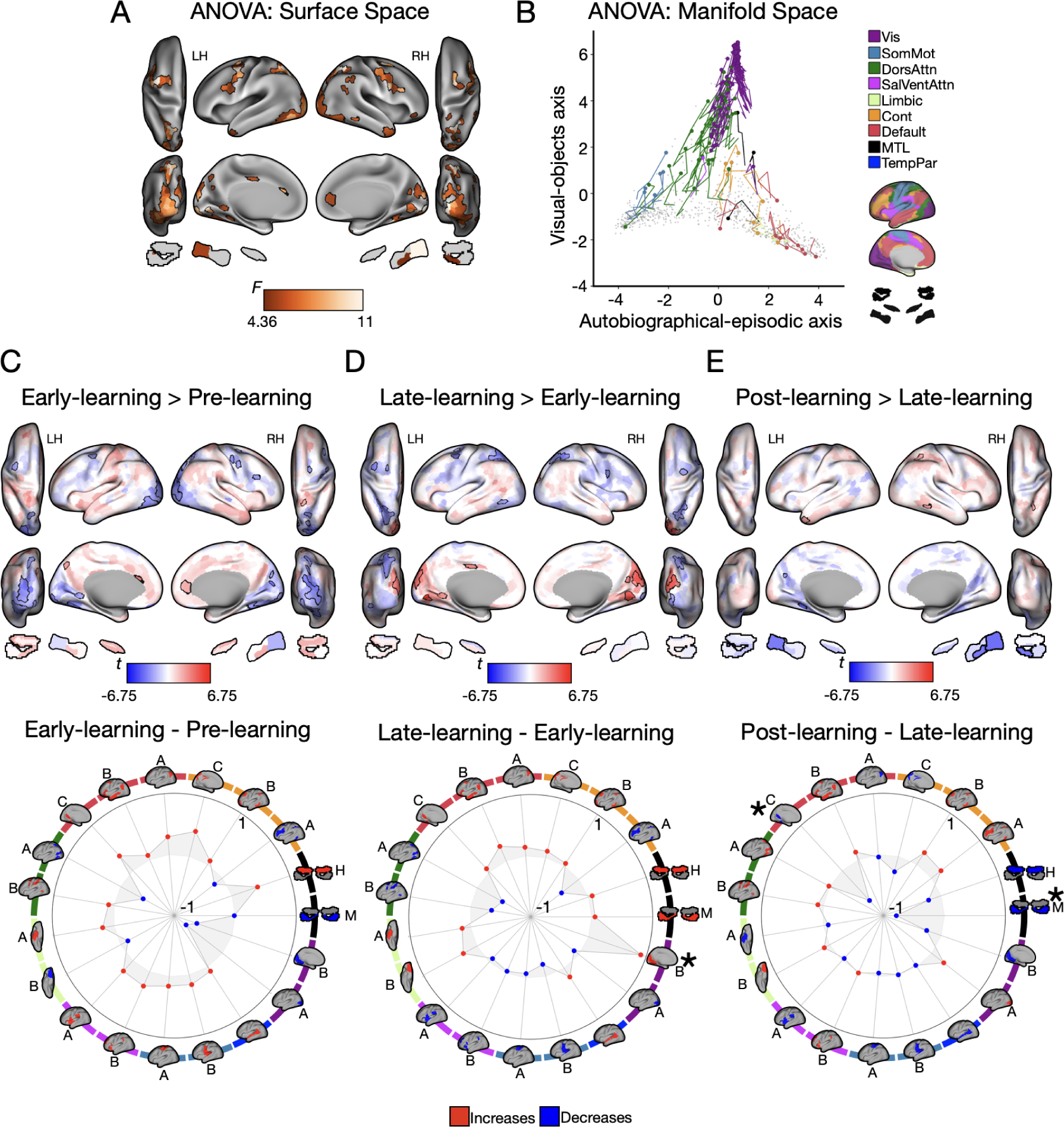
Visual SL is associated with changes along the visual-object axis. (A) Significant regions across across task epochs. (B) Temporal trajectories of regions from A. Each point denotes a region, with significant regions colored by their functional network assignments on the right^55^. Circles denote each regions embedding in Pre-learning, and traces represent trajectories along the axes throughout Early-, Late- and Post-learning. (C-E) Top, Unthresholded pairwise maps. Outlined regions denote significant comparisons. Bottom, Polar plots depicting mean network loading differences for comparisons. The color behind each brain indicates its functional network assignment, and letters depict its subnetwork assignment^55^. Asterisks indicate FDR-corrected significance^56^. Positive (red) values indicate increases (expansion along the axis) and negative (blue) values indicate decreases (contraction along the axis).

To characterize these main effects, for each significant brain region we performed follow-up FDR-corrected^56^ (q < 0.05) paired-samples t-tests between epochs and also aggregated across regions (regardless of their significance) within each functional network^55^. To test for significant changes in loading along the visual-object axis at the onset of learning, we performed an Early-learning > Pre-learning contrast. This contrast revealed contractions within the visual cortex and DAN (outlined in blue; see Fig. 5C top), which could also be observed qualitatively when aggregated at the level of functional network (see Fig. 5C bottom). Note, however, that the network-level changes in loadings between the Pre- and Early-learning epochs were not significant following FDR-correction. (also note that, for the polar plots, we used the 17-network cortical mapping in addition to mapping the hippocampus and MTL to improve spatial precision compared to the 7-network mapping^45,46,55^).

To examine changes from the onset of learning to the end of SL, we performed a contrast of Late-learning > Early-learning. This contrast revealed that regions within the visual cortex, which had originally contracted, now significantly expanded along the visual-object axis (red areas in Fig. 5D top), whereas areas of the DAN contracted further (compare Fig. 5D top with 5C top). However, when extrapolated to the whole-brain functional network level, we found that only the visual network showed significant changes from Early- to Late-learning (see Fig. 5D, bottom).

Although these above neural changes from Early- to Late-learning are interpreted as reflecting an intrinsic process related to SL, it is possible that they are merely attributable to subject fatigue or habituation (i.e., the passage of time). Behavior from the perceptual cover task however, suggests this alternative explanation is unlikely, as participants displayed faster reaction times and greater accuracy from both Early- to Late-Learning, and from Pre- to Post-learning (both *p*s < 0.05). Nevertheless, to further rule out this alternative at the neural level, we examined whether the observed pattern of changes (i.e., expansion of visual cortex regions) also persisted into the final, Post-learning epoch, when SL was interrupted. When we contrasted the Post-learning > Late-learning epochs, we instead found that regions within the MTL — specifically the PRC and ERC — as well as the retrosplenial cortex which did not exhibit significant changes during SL, now modified their connectivity and significantly contracted along the visual-object axis (see Fig. 5E, top). When extrapolated to the network-level, this resulted in significant changes in both the MTL and DMN-C (see Fig. 5E, bottom). This result provides strong support for the interpretation that the neural changes described earlier reflect processes intrinsic to SL rather than mere time-dependent effects.

### Alterations in regional connectivity that underlie changes along the visual-objects axis

While significant changes in the embedding of regions along the visual-objects axis indicate a change to their overall connectivity profile, this does not itself reveal *how* these regions change their connectivity with other areas. To elucidate these changes, we performed seed connectivity analyses using representative regions from left and right hemispheres of the visual network, DAN, and PRC. Together, these regions capture the main patterns of effects observed with respect to the expansions and contractions along the visual-objects axis (see Fig. 6 for seed analyses of the right hemisphere; see Fig. S1 for the left hemisphere). Importantly, we used the same seed regions highlighted in Fig. 4B, allowing us to directly link changes in whole-brain functional connectivity to displacements along the visual-object axis.

**Figure 6.**
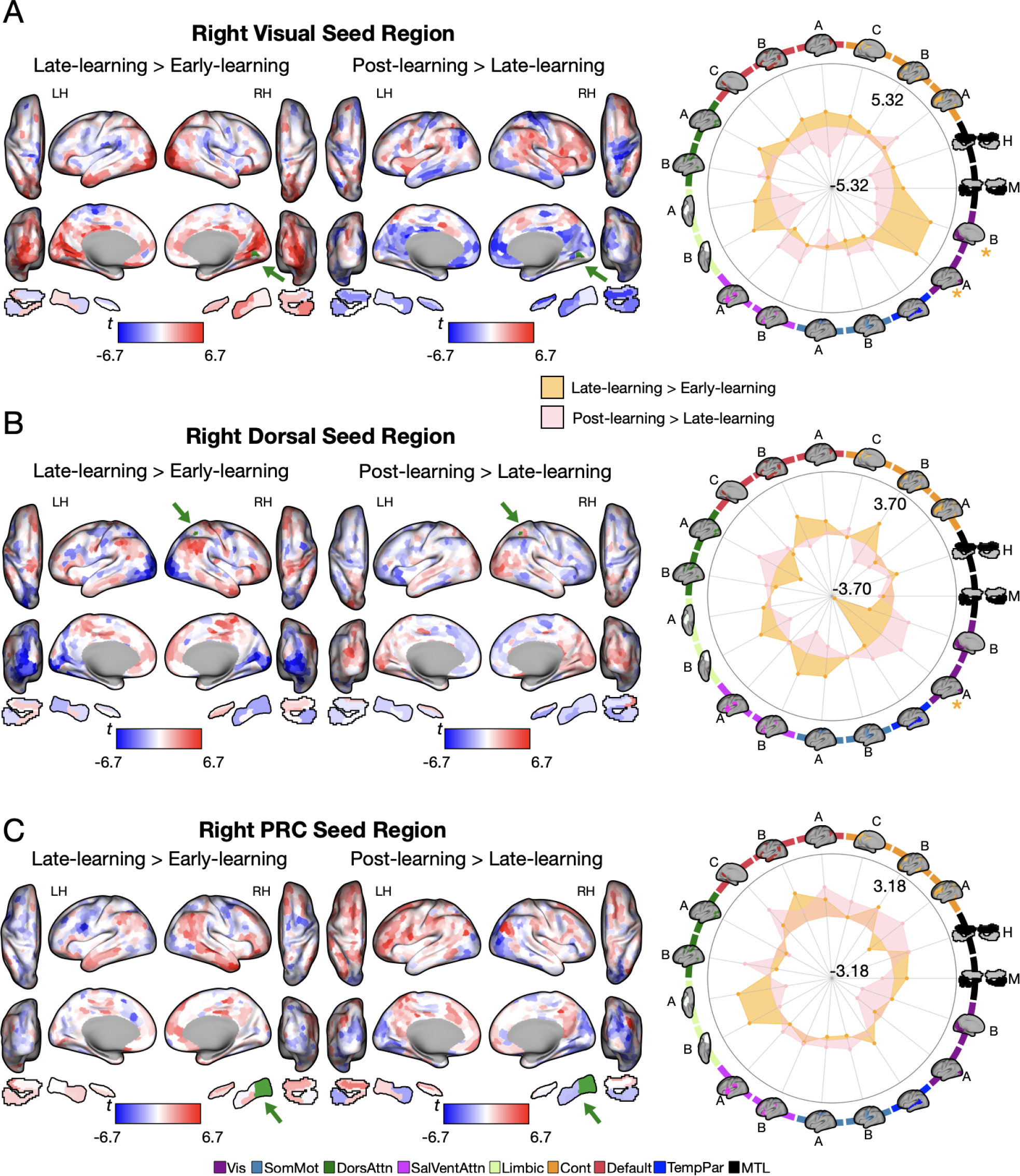
Patterns of connectivity underlying changes along the visual-objects axis throughout visual SL. (A) Connectivity changes of representative seed regions in right visual, DAN, and PRC areas (shown in green, as indicated by green arrow) from Late-learning > Early-learning and Post-learning > Late-learning. Positive (red) values show increases and negative (blue) values show decreases in connectivity. (B) Polar plots show connectivity changes between epochs at the network level (according to the Yeo 17-networks parcellation^55^, as well as MTL regions derived from the Automated Segmentation of Hippocampal Subfields^46^ (ASHS) split into MTL (M) and hippocampus (H)). The color behind each brain indicates its functional network assignment (legend in B), with letters depicting its subnetwork assignment^55^. Asterisks indicate FDR-corrected significance in network changes between epochs.

For each seed region, we contrasted the associated whole-brain connectivity maps for Early- to Late- and Late- to Post-learning with region-wise paired-samples t-tests. In Fig. 6 we display (1) the unthresholded region-wise contrast maps for each seed region to allow for a complete visualization of a given region’s connectivity changes (see Fig. 6 left), and (2) corresponding polar plots, depicting the network-level changes in connectivity for each region (see Fig. 6 right). Together, these plots provide a comprehensive description of the collective changes in connectivity that contribute to the changes in regional embedding along the visual-objects axis.

For the right visual seed region (see Fig. 6A), which exhibited significant expansion along the visual-object axis from Early- to Late-learning (see Fig. 5D), connectivity during SL was primarily characterized by increased covariance with other visual network areas, as well as with regions in the DMN and limbic network. By contrast, from Late- to Post-learning, this connectivity pattern reversed, with decreased covariance in these initial areas and the MTL, and increased covariance with the SalVentAttn network. Although these reversals in connectivity did not result in a statistically significant contraction of this seed region along the visual-object axis during Post-learning, this general region of cortex did tend to exhibit contraction during this Post-learning period (see negative changes in blue in Fig. 5E). Together, these results suggest that, during SL, the visual network’s expansion along the visual-objects axis is mainly driven by increased within-network connectivity; however, once SL is interrupted, these patterns of connectivity appear to reverse, with visual cortex exhibiting increased connectivity with brain networks linked to exogenous attention (SalVentAttn network).

For the right DAN (see Fig. 6B), which exhibited significant *contraction* along the visual-object axis from Early- to Late-learning (see Fig. 5D), we found that its connectivity during SL was most characterized by a decrease in connectivity with other DAN areas and the visual cortex, and an increase in connectivity with higher-order transmodal regions belonging to the DMN B, Control B and SalVentAttn B subnetworks. Conversely, from Late- to Post-learning, when this region significantly *expanded* along the visual-object axis, we observed a stark reversal in connectivity patterns. The DAN region now showed increased connectivity with areas within its own network, as well as with regions of the visual network. These findings suggest that the DAN’s contraction during learning is driven by decreased within-network connectivity and increased between-network connectivity with higher-order brain networks. In contrast, its expansion during Post-learning is characterized by increased within-network connectivity and reinstated connectivity with the visual cortex.

Finally, for the right PRC (see Fig. 6C), which exhibited significant contraction along the visual-object axis from Late- to Post-learning, our seed connectivity maps revealed that, during Post-learning, the PRC decreased connectivity with the visual cortex, while increasing connectivity with large swaths of parietal and prefrontal cortex. The fact that the PRC only significantly changed its embeddings along the visual-object axis once SL was interrupted suggests that it is responsive to prediction errors, and may communicate these errors with the parietal and prefrontal cortex.

Collectively, these seed-based analyses allow us to characterize, at the level of whole-brain functional connectivity, the selective changes in areas along the visual-objects axis both during SL and once the learned associations are violated (during Post-learning). The findings generally support the interpretation that expansions along the visual-object axis reflect, in part, increased within-network connectivity, while contractions indicate increased between-network connectivity.

### The role of the hippocampus in statistical learning

In previous sections, we characterized regional changes along the visual-objects axis across the visual, DAN, and PRC during visual SL, as well as the underlying shifts in functional connectivity driving these changes. While our findings in the PRC suggest its role in processing prediction errors, we observed no significant changes in the embedding of hippocampal regions across our task — a conspicuous absence given prior work implicating hippocampus in SL^17,21^, and evidence for representational change in our earlier analyses^18^ (Fig. 1C). One explanation for this discrepancy may stem from our functional connectivity analysis including 998 cortical but only eight hippocampal regions. Since dimension reduction approaches like PCA aim to explain the largest amount of variance in a relatively small number of orthogonal components, it is not surprising that our strongest loadings (PCs 1-3) were dominated by cortex, and not the hippocampus (see Fig. 3A). It may also be that the changes that occur in the hippocampus are largely internal, and not observable using this higher-level analysis that predominantly focuses on macro-scale changes in connectivity across networks. Nevertheless, as prior work has identified a clear role for the hippocampus in SL, we next sought to directly examine hippocampal connectivity changes during and after learning. To this end, we aggregated seed-based hippocampal connectivity maps for subfields in left and right CA1, CA2/3, DG and subiculum, and created contrast maps to compare Early- to Late-, and Late- to Post-learning. For an overview of our hippocampal-seed analyses, see the Supplementary Extended Materials and Methods.

Our hippocampal-based seed connectivity analysis (see Fig. 7A; for the left hippocampus, see Fig S2) revealed that from Early- to Late-learning, the right hippocampus showed modest qualitative increases in connectivity with all other networks, but specifically with other areas of the MTL, and limbic-A subnetwork (see Figure. 7B). However, once SL was interrupted during Post-learning, these patterns tended to reverse, with the hippocampus most notably decreasing its connectivity with the visual cortex. In addition, the hippocampus also increased connectivity with the contralateral hemisphere and the retrosplenial subdivision of the DMN (i.e., DMN-C subnetwork). While this reversal in connectivity aligns with changes within the PRC and the overall decrease along the visual-objects axis (Fig. 5E), it’s notable that no significant changes were observed in the hippocampus. This, in conjunction with the striking patterns seen in other MTL regions, suggests that the PRC and ERC may play a more crucial role than previously thought in mediating information exchange between the cortex and hippocampus during learning and when the learned associations are violated.

**Figure 7.**
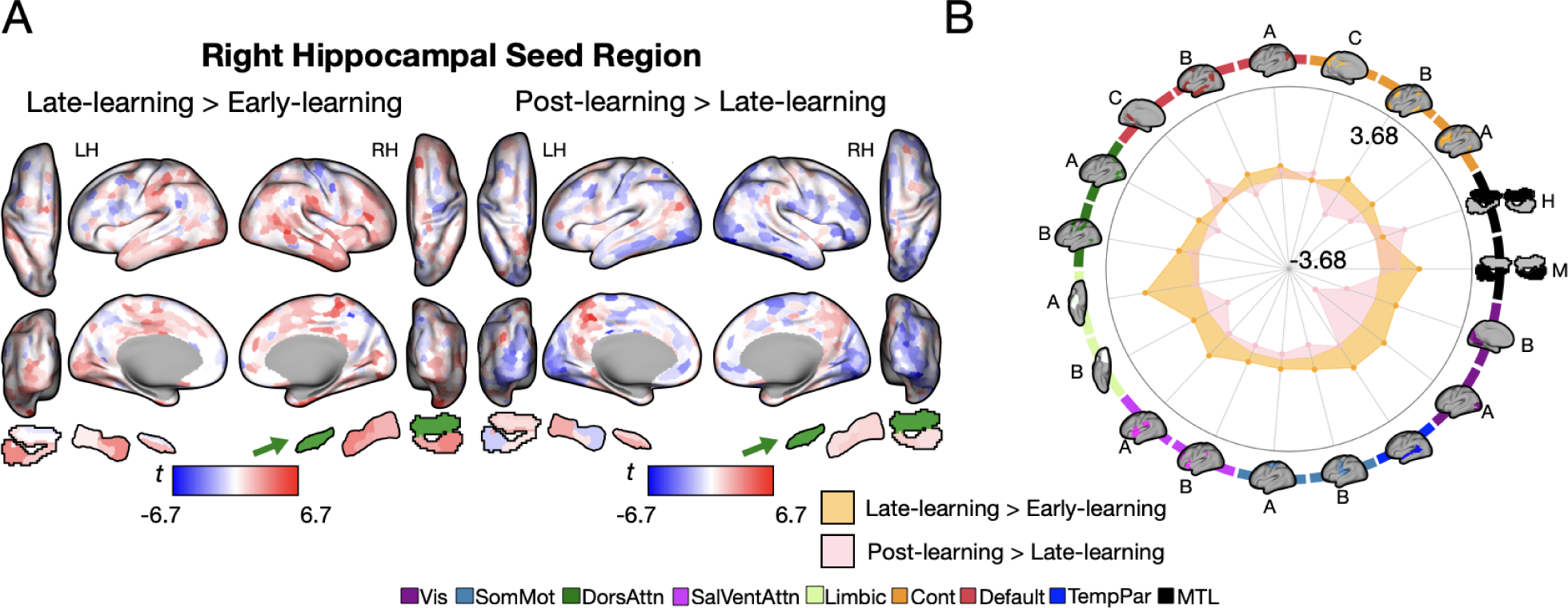
Hippocampal-Based Connectivity Analysis. (A) Connectivity changes for the right hippocampus (in green) from Late-learning > Early-learning and Post-learning > Late-learning. Positive (red) and negative (blue) values show increase and decreases in connectivity, respectively. (B) Polar plots show seed-based changes in connectivity between epochs at the network level (according to the Yeo 17-networks parcellation^55^, as well as the additional MTL regions derived from the Automated Segmentation of Hippocampal Subfields^46^ (ASHS) split into MTL (M) and hippocampus (H)). The color behind each brain indicates its functional network assignment, with letters depicting its constituent subnetwork assignment^55^.

## DISCUSSION

Our study employed dimension reduction techniques to elucidate the neural mechanisms underlying visual SL. By constructing a low-dimensional manifold characterizing the principal axes of whole-brain connectivity, we identified a single dimension — the visual-object axis — along which regions significantly altered their expression during learning and subsequent violation of the learned association. Focused analyses of this axis revealed that during SL, higher-order visual cortex regions expanded, reflecting increased within-network integration and segregation from other networks. Conversely, dorsal attention network (DAN) areas contracted, indicating enhanced integration with other networks and decreased covariance with the visual cortex. Importantly, when learned associations were violated post-learning, we observed a markedly different pattern: MTL regions, previously unaffected during initial learning, exhibited contraction, while the trends in visual and DAN regions either halted or reversed. Collectively, our findings offer new insights into the whole-brain dynamics underlying visual SL. They suggest that, at the macro-scale, SL is predominantly characterized by cortical connectivity changes, whereas the MTL, particularly extra-hippocampal regions like the perirhinal and entorhinal cortices, respond to violations in the learned associations. This suggests a nuanced interplay between cortical and subcortical structures in supporting SL, providing a different perspective on the neural underpinnings of this fundamental cognitive process.

From Early- to Late-learning, we found that regions within the visual network significantly expanded along the visual-objects axis. This expansion was primarily characterized by increased within-network connectivity, likely reflecting a mechanism where the modularity of task-specific networks is increased to insulate encoded associations from additional change^57,58^ and thus enhance the robustness of learning. In contrast, the DAN’s contraction along this axis was driven by decreased connectivity with visual cortex, and increased connectivity with areas of transmodal cortex (Control and SalVentAttn networks), consistent with prior work showing reduced connectivity between visual and higher-order association regions as SL takes place^8^. This suggests that as regularities are encoded, cross-talk between task-specific sensory areas and those involved in attentional control becomes less important for learning.

A novel aspect of our task involved the inclusion of SL blocks that were book-ended by unstructured runs, allowing us to investigate changes in whole-brain activity when learned associations were disrupted (in Post-learning). In these post-learning blocks, we observed widespread reversals in several of the whole-brain seed connectivity patterns. Indeed, regions that had changed their covariance with visual regions and DAN during learning showed opposite patterns after SL was interrupted. Specifically, the visual cortex, which had become more modular during learning, now integrated with higher-order attention networks. Similarly, while the DAN became less modular during SL, it now increased within-network connectivity and connectivity with the visual network. These reversals in connectivity suggest the brain’s reorganization during SL is partially undone when SL is disrupted, possibly reflecting a response to a prediction error signal (see more on this below).

One noteworthy finding was that the hippocampus, despite showing theory-consistent representational change as a result of SL, did not exhibit significant connectivity changes during or after learning. However, instead we found that extra-hippocampal MTL regions, which did not initially show learning effects — the PRC and ERC — now significantly contracted along the visual-objects axis in Post-learning. These contractions reflected increased connectivity with the DAN, DMN, and higher-order control networks, and decreased covariance with the visual cortex. Current frameworks highlight the PRC as a specialized region positioned at the apex of the ventral visual stream, supporting object recognition^17,59,60^, and the ERC as a bridge between the ventral visual pathway and hippocampus, via (in part) PRC^59,61^. The PRC and ERC’s contraction likely reflects a response to prediction error, where regions recouple with higher-order cortical networks to reorient attention to incoming sensory information. Prediction error signals evoked in the hippocampus^62,63^, propagate to cortical regions implicated in perception and attention to update existing schemas^64,65^, propagate to cortical regions implicated in perception and attention to update existing schemas^66^. Our findings not only extend this theoretical view of SL, but provide a novel perspective of how such prediction errors may be distributed across the cortex. Indeed, while prior work has focused on the role of the PRC and ERC in more explicit forms of learning^6,7,67^, our results suggest a key role for these regions in detecting violations in implicit learning, perhaps precipitating the reversals observed in our cortical seed regions. Similarly, while prior work has largely implicated the hippocampus in prediction error, our findings indicate that MTL regions may play a larger role in controlling the flow of information than previously assumed.

The differential role of cortical and MTL networks in our main analysis (see Fig. 5) provides a nuanced understanding of whether SL operates primarily through cortical or MTL mechanisms. Specifically, our results highlight that interactions within and between these regions dynamically update during SL, leading to different network configurations depending on the phase of the task. Both cortical and MTL circuits appear to learn in parallel: the hippocampus undergoes local representational change to encode paired stimuli^16–18^, while sensory cortical circuits become more modular, disconnecting from higher-order cognitive networks. However, when learned regularities are disrupted, the MTL detects this mismatch and reintegrates with these cognitive networks, while decreasing connectivity with the visual cortex. Importantly, our current findings indicate that these macro-scale changes in the landscape of whole-brain activity occur largely independent of the hippocampus, suggesting a prominent role for other MTL regions, like PRC and EC. This highlights the complexity of SL mechanisms, where cortical and MTL regions may contribute in distinct yet complementary ways.

While the current study offers unique insights into the whole-brain changes in functional connectivity associated with visual SL, there remain several important avenues for future research. First, while our findings point to distinct patterns of connectivity change associated with learning and the later violation of that learning, future work could directly track the behavioral correlates of these neural changes on a more moment-by-moment basis. Second, it remains underexplored how the neural changes observed here may relate to other forms of implicit learning (e.g., auditory, spatial SL, error-based learning), as well as to more general distinctions made between implicit versus explicit forms of learning^68^. For example, in the sensorimotor control literature, implicit learning is thought to be cerebellar-dependent, and yet it is unknown whether SL also relies on cerebellum, which contains similar functional networks to cortex^69^. Third, given its established role in SL and prediction^70^, it was surprising to us that the hippocampus did not significantly change its embedding along the visual-objects axis during any phases of the task. In future work, it might be advantageous to construct a hippocampus-centric manifold to increase the sensitivity in detecting changes in its patterns of covariance; alternatively, our findings and whole-brain approach may highlight the relatively greater importance of other regions (outside the hippocampus) in guiding SL and encoding violations of learned associations.

In summary, our work provides a unique whole-brain perspective explaining the connectivity changes that occur during SL. The findings indicate that while cortical and MTL regions are involved in SL, this may occur via different mechanisms and at during different time periods, where cortical changes subserve learning through repetition, while MTL handles reorganization to allow for updating in response to unexpected deviations from prior learning.

## MATERIALS AND METHODS

We re-analyzed data^18^ of 33 healthy individuals (21 female) between the ages of 18-35 years old, completing a visual SL task. Eight of the original dataset’s 41 participants were excluded either because they completed fewer task runs than the majority of the sample (n = 7), or because there were issues with the quality of hippocampal or cortical segmentations (n = 1).

Participants completed eight experimental runs, where in each, 16 abstract images were presented one at a time on screen, each repeated five times, for a total of 80 image presentations. In all runs, the participants’ unrelated cover task was to identify rare (10% of trials) instances of a small, faded grey square being overlaid on the images, by pressing a button. In the first experimental run (Pre-learning), the images were presented in a completely random, unstructured order. Following this run, participants completed six consecutive experimental runs (80 image presentations each, 480 total), in which unbeknownst to them, the 16 images were covertly combined into 8 pairs. This pair structure was embedded into the image sequence, such that a given image (A) was always followed by its pair-mate (B). The first and last of these structured runs were used as Early- and Late-learning epochs respectively, for the present analysis. Finally, after SL, participants performed an additional Post-learning run, where they again viewed the 80 images in a completely random order (similar to Pre-learning).

For each participant and epoch, we obtained region-wise timeseries data by taking the mean BOLD activity for each of the 998 cortical regions pre-defined by the Schaefer-1000 parcellation^45^ (two regions were removed due to small parcel size) and 14 MTL regions, derived in an individual-specific way using ASHS software^46^. Across the superset of these 1012 regions, we created centered functional connectivity matrices for each epoch, transforming them into cosine similarity affinity matrices after applying row-wise proportional thresholding. Next, we used PCA on affinity matrices to construct connectivity manifolds^35,36,50^ and aligned all PCs to a group-average template Pre-learning manifold using Procrustes transformation.

Guided by prior work^40^ we uncovered task-related variations within our top three PCs, by quantifying each subject’s variability in PC loadings (min-max spread) across our task epochs. For each PC, we performed a rmANOVA on subjects’ difference scores, and corrected for multiple comparisons using FDR correction^56^ (q < 0.05). This analysis revealed a significant effect of epoch along a single neural dimension, the visual-objects axis. To evaluate learning-related changes in whole-brain functional connectivity over time, we computed each brain region’s loadings along the visual-objects axis and assessed region-wise differences in loadings across epochs using rmANOVAs and follow-up paired t-tests. To explore alterations in whole-brain connectivity underlying changes along this axis, we performed seed connectivity contrasts on representative seed regions (belonging to the visual network, DAN, and PRC) that demonstrated representative effects both during and after SL. To examine learning-related changes relative to the hippocampus, we also performed similar seed connectivity contrasts for hippocampal regions to examine changes both during and after SL. For a comprehensive overview of the methods, please refer to the Supplementary Extended Materials and Methods.

## Code availability

Imaging data were preprocessed using fmriPrep, which is open source and freely available. Our analysis is available at https://github.com/thelamplab/visualSLmanifolds.

