## Supporting Information for "Visual statistical learning alters low-dimensional cortical architecture"

### EXTENDED METHODS AND MATERIALS

**Participants**. Our study performed a re-analysis on data collected from 33 healthy individuals (21 female) between the ages of 18-35 years old (1). All participants reported having normal visual acuity (either uncorrected or corrected to normal) and good colour vision. All participants gave informed consent to a protocol approved by the Yale Institutional Review Board (IRB) and received $20 USD per hour as compensation for their time. Eight participants were excluded from the original dataset's 41 participants due to either missing functional runs (n = 7) or issues in the quality of hippocampal and cortical segmentations in the present data analysis (n = 1).

**Statistical learning task.** To investigate changes in whole-brain functional connectivity during statistical learning (SL), we analyzed human fMRI data from a visual SL task, where abstract images were presented to subjects one at a time on screen (1). The task stimuli included eight unique image pairs, varying in similarity from having no discernible shared features to being nearly indistinguishable from each other (see Main Paper Fig. 1A). The task was composed of eight functional runs, each lasting 304.5 seconds (203 TRs), where images were presented for 1 second each. These functional runs took two different forms: structured or random. In the structured runs, unbeknownst to the participant, the sequence of images presented to the participant had the eight image pairs (e.g., AB, CD) embedded, such that when a given image (e.g., A) was presented, it was always followed by its pair-mate (e.g., B). A given pair in the structured runs was never presented twice in a row. In visual SL tasks, participants come to implicitly associate images that are paired together over time, as established in prior behavioural and neural data (2-4). Participants completed six of these structured runs in a row.

These six structured runs were bookended by two unstructured runs, where images were presented in a random order (i.e., unpaired), thereby allowing us to establish the characteristic patterns of whole-brain connectivity that manifest prior to learning (Pre-learning run), as well as after learning had occurred and the paired sequence structure was interrupted (Post-learning run). During both structured and random runs, a small, partially transparent gray patch was overlaid on 10% of images (see Main Paper Fig. 1B). The participants were instructed to perform a cover task of pressing a button on a hand-held box when they noticed the gray patch. This explicit, secondary task was entirely orthogonal to the pair structure embedded in the SL task, and simply served to maintain attention on the images throughout the task.

**MRI acquisition and preprocessing.** All data collected from the prior research (1) was scanned using a 3T Siemens Prisma scanner with a 64-channel head coil at the Yale Magnetic Resonance Research Center. For each participant, eight functional runs were collected with a multiband echo planar imaging (EPI) sequence (TR = 1500 ms; TE = 3.26 ms; voxel size = 1.5mm isotropic; FA = 71; multiband factor = 6), yielding 90 axial slices. Each run contained 203 volumes. Results included in this manuscript come from preprocessing performed using fMRIPrep 20.1.2 (5-6; RRID: SCR_016216), which is based on Nipype 1.5.1 (7-8; RRID: SCR_002502). Many internal operations of fMRIPrep use Nilearn 0.6.2 (9, RRID:SCR_001362), mostly within the functional processing workflow. For more details of the pipeline, see the section corresponding to workflows in fMRIPrep's documentation.

**Anatomical data preprocessing.** A total of 1 T1-weighted (T1w) images were found within the input BIDS dataset. The T1-weighted (T1w) image was corrected for intensity non-uniformity (INU) with N4BiasFieldCorrection (10), distributed with ANTs 2.2.0 (11, RRID:SCR_004757), and used as T1w-reference throughout the workflow. The T1w-reference was then skull-stripped with a Nipype implementation of the antsBrainExtraction.sh workflow (from ANTs), using OASIS30ANTs as target template. Brain tissue segmentation of cerebrospinal fluid (CSF), white-matter (WM) and gray-matter (GM) was performed on the brain-extracted T1w using fast (FSL 5.0.9, RRID:SCR_002823, 12). Brain surfaces were reconstructed using recon-all (FreeSurfer 6.0.1, RRID:SCR_001847, 13), and the brain mask estimated previously was refined with a custom variation of the method to reconcile ANTs-derived and FreeSurfer-derived segmentations of the cortical gray-matter of Mindboggle (RRID:SCR_002438, 14). Volume-based spatial normalization to two standard spaces (MNI152NLin6Asym, MNI152NLin2009cAsym) was performed through nonlinear registration with antsRegistration (ANTs 2.2.0), using brain-extracted versions of both T1w reference and the T1w template. The following templates were selected for spatial normalization: FSL's MNI ICBM 152 non-linear 6th Generation Asymmetric Average Brain Stereotaxic Registration Model [15, RRID:SCR_002823; TemplateFlow ID: MNI152NLin6Asym], ICBM 152 Nonlinear Asymmetrical template version 2009c [16, RRID:SCR_008796; TemplateFlow ID: MNI152NLin2009cAsym].

**Functional data preprocessing.** For each of the 8 BOLD runs found per subject (across all tasks and sessions), the following preprocessing was performed. First, a reference volume and its skull-stripped version were generated using a custom methodology of fMRIPrep. Head-motion parameters with respect to the BOLD reference (transformation matrices, and six corresponding rotation and translation parameters) are estimated before any spatiotemporal filtering using mcflirt (FSL 5.0.9, 17). BOLD runs were slice-time corrected using 3dTshift from AFNI 20160207 (18, RRID:SCR_005927). A B0-nonuniformity map (or fieldmap) was estimated based on two (or more) echo-planar imaging (EPI) references with opposing phase-encoding directions, with 3dQwarp (18; AFNI 20160207). Based on the estimated susceptibility distortion, a corrected EPI (echo-planar imaging) reference was calculated for a more accurate co-registration with the anatomical reference. The BOLD reference was then co-registered to the T1w reference using bbregister (FreeSurfer) which implements boundary-based registration (19). Co-registration was configured with six degrees of freedom. The BOLD time-series were resampled onto the following surfaces (FreeSurfer reconstruction nomenclature): fsaverage5, fsaverage. The BOLD time-series (including slice-timing correction when applied) were resampled onto their original, native space by applying a single, composite transform to correct for head-motion and susceptibility distortions. These resampled BOLD time-series will be referred to as preprocessed BOLD in original space, or just preprocessed BOLD. The BOLD time-series were resampled into standard space, generating a preprocessed BOLD run in MNI152NLin6Asym space. First, a reference volume and its skull-stripped version were generated using a custom methodology of fMRIPrep. Grayordinates files (20) containing 91k samples were also generated using the highest-resolution fsaverage as intermediate standardized surface space. Automatic removal of motion artifacts using independent component analysis (ICA-AROMA, 21) was performed on the preprocessed BOLD on MNI space time-series after removal of non-steady state volumes and spatial smoothing with an isotropic, Gaussian kernel of 6mm FWHM (full-width half-maximum). Corresponding "non-aggresively" denoised runs were produced after such smoothing. Additionally, the "aggressive" noise-regressors were collected and placed in the corresponding confounds file. Several confounding time-series were calculated based on the preprocessed BOLD: framewise displacement (FD), DVARS and three region-wise global signals. FD was computed using two formulations following Power (absolute sum of relative motions, 22) and Jenkinson (relative root mean square displacement between affines, 17). FD and DVARS are calculated for each functional run, both using their implementations in Nipype (following the definitions by 22). The three global signals are extracted within the CSF, the WM, and the whole-brain masks.

Additionally, a set of physiological regressors were extracted to allow for component-based noise correction (CompCor, 23). Principal components are estimated after high-pass filtering the preprocessed BOLD time-series (using a discrete cosine filter with 128s cut-off) for the two CompCor variants: temporal (tCompCor) and anatomical (aCompCor). tCompCor components are then calculated from the top 5% variable voxels within a mask covering the subcortical regions. This subcortical mask is obtained by heavily eroding the brain mask, which ensures it does not include cortical GM regions. For aCompCor, components are calculated within the intersection of the aforementioned mask and the union of CSF and WM masks calculated in T1w space, after their projection to the native space of each functional run (using the inverse BOLD-to-T1w transformation). Components are also calculated separately within the WM and CSF masks. For each CompCor decomposition, the k components with the largest singular values are retained, such that the retained components' time series are sufficient to explain 50 percent of variance across the nuisance mask (CSF, WM, combined, or temporal). The remaining components are dropped from consideration. The head-motion estimates calculated in the correction step were also placed within the corresponding confounds file. The confound time series derived from head motion estimates and global signals were expanded with the inclusion of temporal derivatives and quadratic terms for each (24). Frames that exceeded a threshold of 0.5 mm FD or 1.5 standardised DVARS were annotated as motion outliers. All resamplings can be performed with a single interpolation step by composing all the pertinent transformations (i.e. head-motion transform matrices, susceptibility distortion correction when available, and co-registrations to anatomical and output spaces). Gridded (volumetric) resamplings were performed using antsApplyTransforms (ANTs), configured with Lanczos interpolation to minimize the smoothing effects of other kernels (25). Non-gridded (surface) resamplings were performed using mri_vol2surf (FreeSurfer).

**Reanalysis of the nonmonotonic plasticity hypothesis.** Evidence supporting visual SL in the task comes from a re-analysis from the original paper of this dataset (1), subsetted for only the subjects included in our analysis. Here, we tested for evidence of representational change that is in line with predictions from the nonmonotonic plasticity hypothesis (NMPH; 26-29). That is, representational change is expected to follow a cubic function based on image pair similarity, where pairmates with moderate levels of similarity will cause a dip towards neural differentiation, as opposed to images with no visual similarity where no change is expected to occur, and high-levels of similarity where integration is expected. We tested for this predicted pattern in the hippocampus, as well as in the dentate gyrus (DG), CA1, and CA2/3 as per the prior paper (1).

As a brief overview of the representational change analysis used in the original paper, a general linear model was fit for each of the 16 images for the two random runs (here, referred to as the Pre- and Post-learning epochs). The parameter estimates from each of the unique images were extracted from the voxels within each region of interest (CA1, CA2/3, DG, and hippocampus), and vectorized. For each image pair, we then computed a Pearson correlation between the two vectors representing each image in the pair, yielding eight distinct representational similarity values in both Pre- and Post-learning epochs. For each image pair, these Pre-learning values were then subtracted from Post-learning scores to examine how an image pair's representation changed over the course of visual SL (i.e., from Early- to Late-learning). Here, positive scores indicate integration (i.e. an increase in overlap), while negative scores relate to differentiation (i.e. a decrease in overlap) between the representations of the two images in a given pair. We then tested whether these representational change scores were consistent with the learning-related change predicted by the NMPH. This hypothesis was quantified by fitting a theory-constrained cubic model to the data from all but one held-out participant, then testing how well this model predicted the held-out participant's data. This analysis was completed for each participant in the sample, and the resulting correlations between each participant's observed values, and their model-predicted values were submitted to bootstrap-resampled t-tests. For a full overview of the methodology, please refer to the original paper (1).

**Cortical-subcortical region timeseries extraction.** For each participant and each of our task epochs (Pre-, Early-, Late-, and Post-learning), we extracted the mean blood oxygen level-dependent (BOLD) timeseries for each of the 998 of the 1000 cortical regions pre-defined by the Schaefer 1000 parcellation (30) and each of the 14 medial temporal lobe (MTL) regions and hippocampal subfields pre-defined using the individual-specific parcellations derived from automated segmentations of hippocampal subfields (ASHS) software (31) with the Princeton Atlas (32). Two cortical regions in the Schaefer 1000 parcellation were removed due to their small parcel size. The choice of the Schaefer-1000 parcellation as our method of cortical brain parcellation was based on prior work (33), which provides a balance of both detailed spatial data while accounting for computational feasibility across participants in our analysis. The choice of including hippocampal areas was because of the prominent role of the hippocampus in SL (34-41), and we opted to use individual-specific hippocampal parcellations from ASHS to account for anatomical individual differences within MTL structures. Our timeseries data was denoised using a combination of our confound regressors from our initial preprocessing using fMRIprep, alongside discrete cosine regressions (with a high-pass filter threshold of 128s) generated by fMRIprep. Furthermore, a low pass filtering process using a Butterworth filter with a cut-off point at 100s was also applied, and implemented using Nilearn. Finally, cortical and subcortical timeseries data were z-scored within each region, and combined to create a comprehensive timeseries dataset, including both MTL and cortical activity.

**Functional Connectivity Estimation and Centering.** For each participant, we extracted regional timeseries data for four equal-length task epochs (each containing 203 imaging volumes). The Pre-learning run (i.e., the first random run), where images were presented to participants in an unstructured order, was used as a baseline. The Early- and Late-learning runs consisted of the first and last structured runs in the task, where images were presented in an order that contained an embedded pair structure, such that a given image was always followed by its pairmate. The first and last structured runs were chosen in order to capture a timepoint very early, and a timepoint very late in implicit learning. Finally, we used the final, random run as our measure of Post-learning. For each of our task epochs (Pre-, Early-, Late-, and Post-learning), we generated functional connectivity matrices by calculating the region-wise covariance matrix using the Ledoit-Wolf estimator (42).

As demonstrated in prior work (33, 43), the influence of subject-level clustering on task-based changes in functional connectivity can hinder the analysis of group-level learning-related changes in connectivity. To uncover differences in task-related structure and remove subject-level differences, we used a technique previously described in prior work (44-45), to center the functional connectivity matrices, which leverages the natural Riemannian geometry of the covariance matrix space. In brief, our process involved making adjustments to the covariance matrices of each participant so they all shared a common mean, allowing us to eliminate the variations in functional connectivity that are specific to individual participants (see Main Paper Fig. 2B). For details of this centering approach, as well as its benefits, see (45).

**Dimension reduction and manifold construction.** The following steps were used to derive connectivity manifolds from our previously centered functional connectivity matrices. First, in line with previous research (33, 46-47), we applied row-wise thresholding to retain the top 10% of connections in each row of our matrices to remove any weak or spurious connections. Next, to characterize the similarity between connectivity profiles of different brain regions, we computed cosine similarity between rows in our matrices to create an affinity matrix. We then used Principal Components Analysis (PCA) to extract a set of principal components (PCs) that provide a low-dimensional representation of the connectivity structure of our data.

To investigate changes in participants' functional network architectures across visual SL, we constructed a template manifold from a group-averaged Pre-learning connectivity matrix. This template was calculated from the geometric mean across all participants' centered Pre-learning connectivity matrices. Subsequently, all individual manifolds (33 participants by 4 epochs; 132 total manifolds) were aligned to this template manifold using Procrustes transformation (see Main Paper Fig. 2A). This approach allowed us to examine learning-specific deviations away from the template manifold throughout the task in a common low-dimensional neural manifold space, thus enhancing the sensitivity of our analyses.

**World-cloud analysis.** The loadings from our top three PCs were interpreted using the Neurosynth Decoder (48). This tool employs an automatic meta-analysis and text-mining technique to take specific activations found in different whole-brain maps (in our case, the map for each PC loading; see Main Paper Fig. 3A) to identify keywords that are commonly found in neuroimaging studies with similar maps. It uses this method to generate a collection of keywords that are associated with a given brain mask, along with their correlation values. Our analysis identified the ten most significant positive and negative correlations for each PC's map. We then transformed these lists into word clouds, where the size and color of each term reflected its correlation's direction and strength (see Main Paper Fig. 3E). Importantly, we omitted all terms related to neuroanatomy (such as "prefrontal") and any repeated terms (for instance "object" and "objects") from our final selection.

**Investigating the spread of the top three principle components.** Guided by prior work (46), we investigated task-related variation within our top three PCs by quantifying the variability of each subject's PC loadings across the four task epochs (Pre-, Early-, Late-, and Post-learning). This approach aimed to characterize the range of whole-brain connectivity changes throughout the SL task, and identify which PCs, if any, exhibited significant shifts across these epochs of interest. For each PC, we compared the loadings for each region along this axis, and found the minimum and maximum loading for each participant. We then subtracted the minimum from the maximum to derive a measure of the min-max spread for each participant across each of our task epochs. With these difference score data (i.e. one min-max score for each participant and epoch), we performed a repeated measures ANOVA (rmANOVA) across task epochs for each PC. We corrected each rmANOVA for multiple comparisons using false discovery-rate (FDR) correction at $q < 0.05$ (49). The results of each of these rmANOVAs revealed which neural dimensions were most affected as a function of task epoch, allowing us to discern which dimension to further investigate changes in regional brain activity.

**Examining changes along the visual-objects axis during statistical learning.** In our main analysis, we compared each region's loadings on the visual-objects axis (PC1) during Pre-, Early-, Late- and Post-learning epochs by performing an rmANOVA for each of the 1012 regions in our subcortical-cortical manifold. In addition to the rmANOVAs, we also performed post hoc paired t-tests on significant regions between individual epoch contrasts. These additional t-tests served to further investigate the specific differences from Pre- to Early-learning, Early- to Late-learning, and Late- to Post-learning. Both the ANOVAs and t-tests were corrected for multiple comparisons using FDR-correction ($q < 0.05$; 49).

**Seed Connectivity Analysis.** To explore the underlying changes in functional connectivity that lead to alterations along the visual-objects axis (PC1), we performed seed connectivity contrasts between different task epochs. To this end, we selected seed regions whose loadings on the visual-objects axis were modulated based on task epoch, which included regions in both left and right visual network, DAN and PRC. Together, the patterns of connectivity changes associated with these regions help to describe the underlying shifts in connectivity that result in the key changes in loadings along the axis that were observed in our main analysis.

For each seed region, we created functional connectivity maps for each participant and epoch of interest, and computed region-wise paired t-tests for both Late-learning > Early-learning and Post-learning > Late-learning comparisons. For each comparison, we opted to show the unthresholded t-maps, to better visualize and compare the multivariate changes in connectivity that drive the observed changes along the visual-objects axis. In addition to these t-maps, we constructed polar plots depicting the specific changes in t-values at the network level, by averaging the correlation values for all areas within each network, and then averaging across participants (50). Note that this analyses are mainly intended to provide a visual characterization and interpretation of the connectivity changes of each seed region from our main analysis.

**Hippocampal-Driven Analysis.** In order to characterize the specific changes in functional connectivity between the hippocampus and cortex during our task, we performed an additional analysis, contrasting left and right hippocampal seed connectivity maps across our task epochs. We created these left and right hippocampal seed maps by computing functional connectivity maps for bilateral CA1, CA2/3, DG, and subiculum seed regions, and then averaging the correlation values from each of the connectivity maps within the set of left and right subregions, respectively. By using hippocampal-based maps, we aimed to examine how the hippocampus specifically interacts with other cortical areas throughout different task epochs, a feature that is predicted from the previous literature (51-52), but not apparent from our main analysis. Similar to our main seed connectivity analysis, we also constructed polar plots for each hippocampal seed region to better depict the specific changes in correlations at the network level.

### SI FIGURES

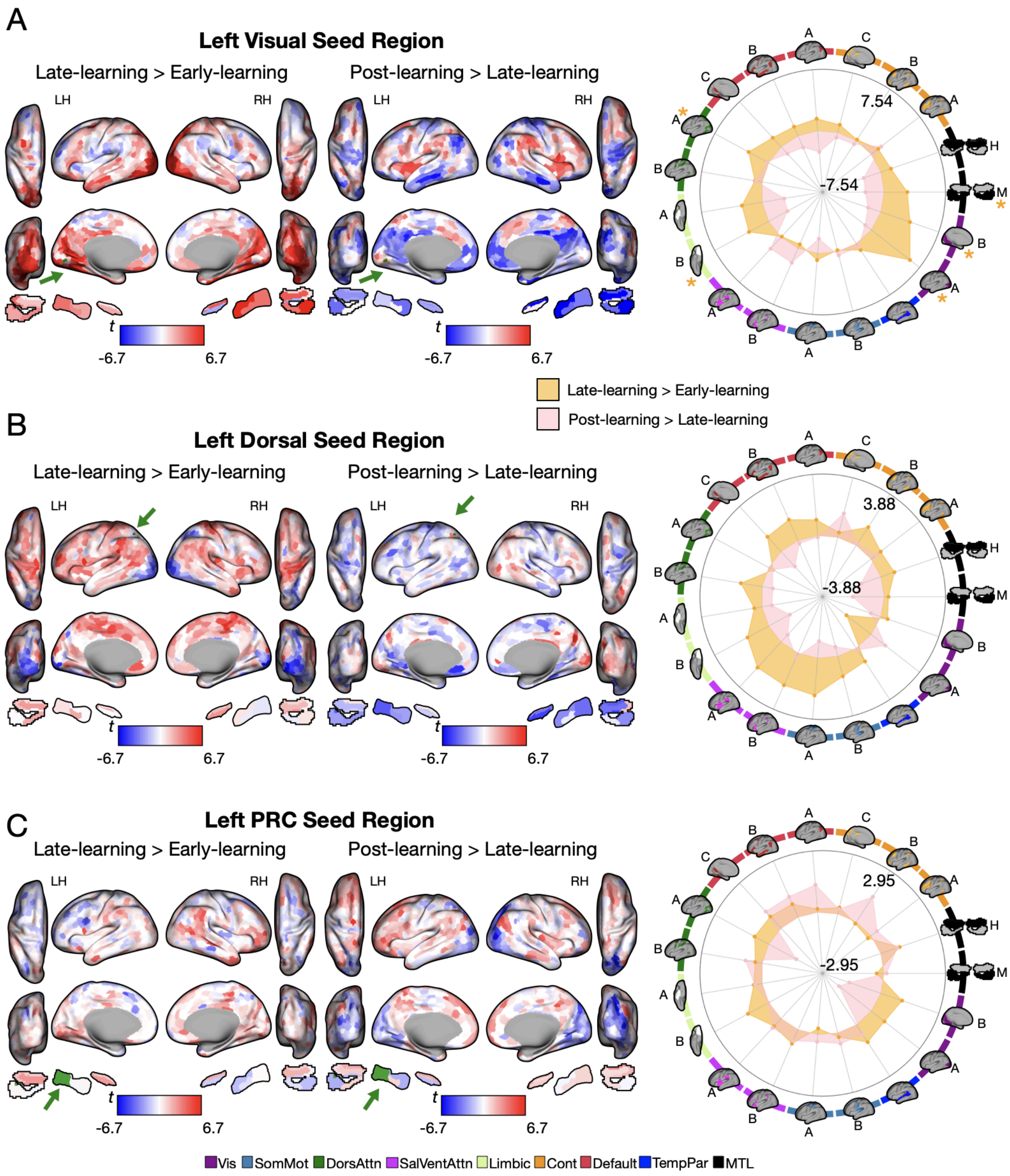


**Figure S1. Patterns of connectivity underlying changes along the visual-objects axis throughout visual SL.** (A) Connectivity changes of representative seed regions in left visual, DAN, and PRC areas (shown in green, as indicated by green arrow) from Late-learning >

Early-learning and Post-learning > Late-learning. Positive (red) values show increases and negative (blue) values show decreases in loading across along the visual-objects axis. (B) Polar plots show seed-based changes in connectivity between epochs at the network level (according to the Yeo 17-networks parcellation (30), as well as the additional MTL regions derived from the Automated Segmentation of Hippocampal Subfields (ASHS; 31) split into MTL (M) and hippocampus (H)). The color behind each brain indicates its functional network assignment (legend in B), with letters depicting its constituent subnetwork assignment (50). Asterisks indicate FDR-corrected significance in network changes between epochs.

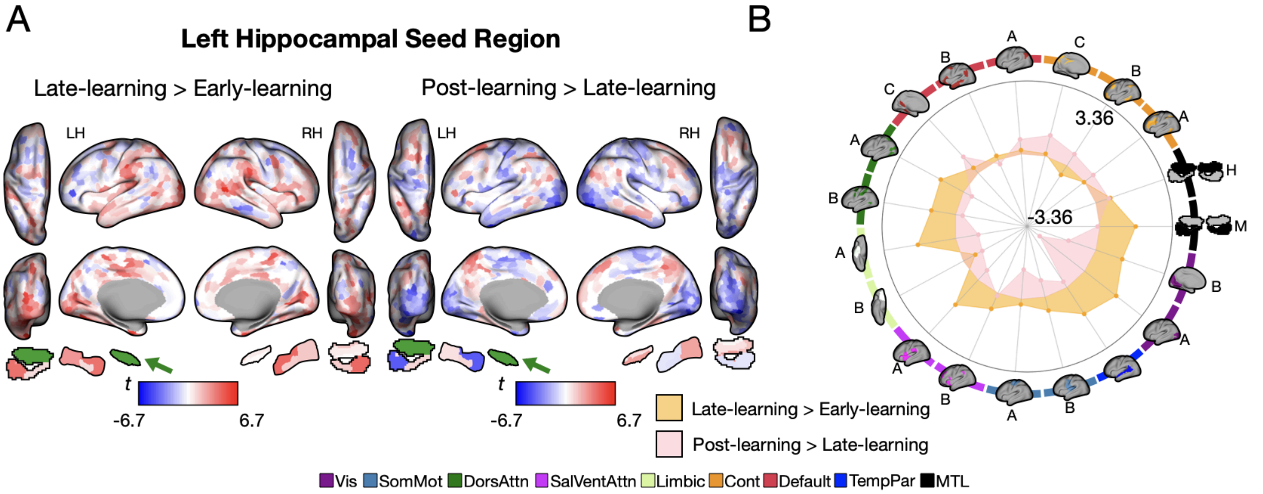


**Figure S2. Hippocampal-Based Connectivity Analysis.** (A) Connectivity changes for the left hippocampus (in green) from Late-learning > Early-learning and Post-learning > Late-learning. Positive (red) and negative (blue) values show increase and decreases in connectivity, respectively. (B) Polar plots show seed-based changes in connectivity between epochs at the network level (according to the Yeo 17-networks parcellation (50), as well as the additional MTL regions derived from the Automated Segmentation of Hippocampal Subfields (ASHS; 31) split into MTL (M) and hippocampus (H)). The color behind each brain indicates its functional network assignment, with letters depicting its constituent subnetwork assignment (50).

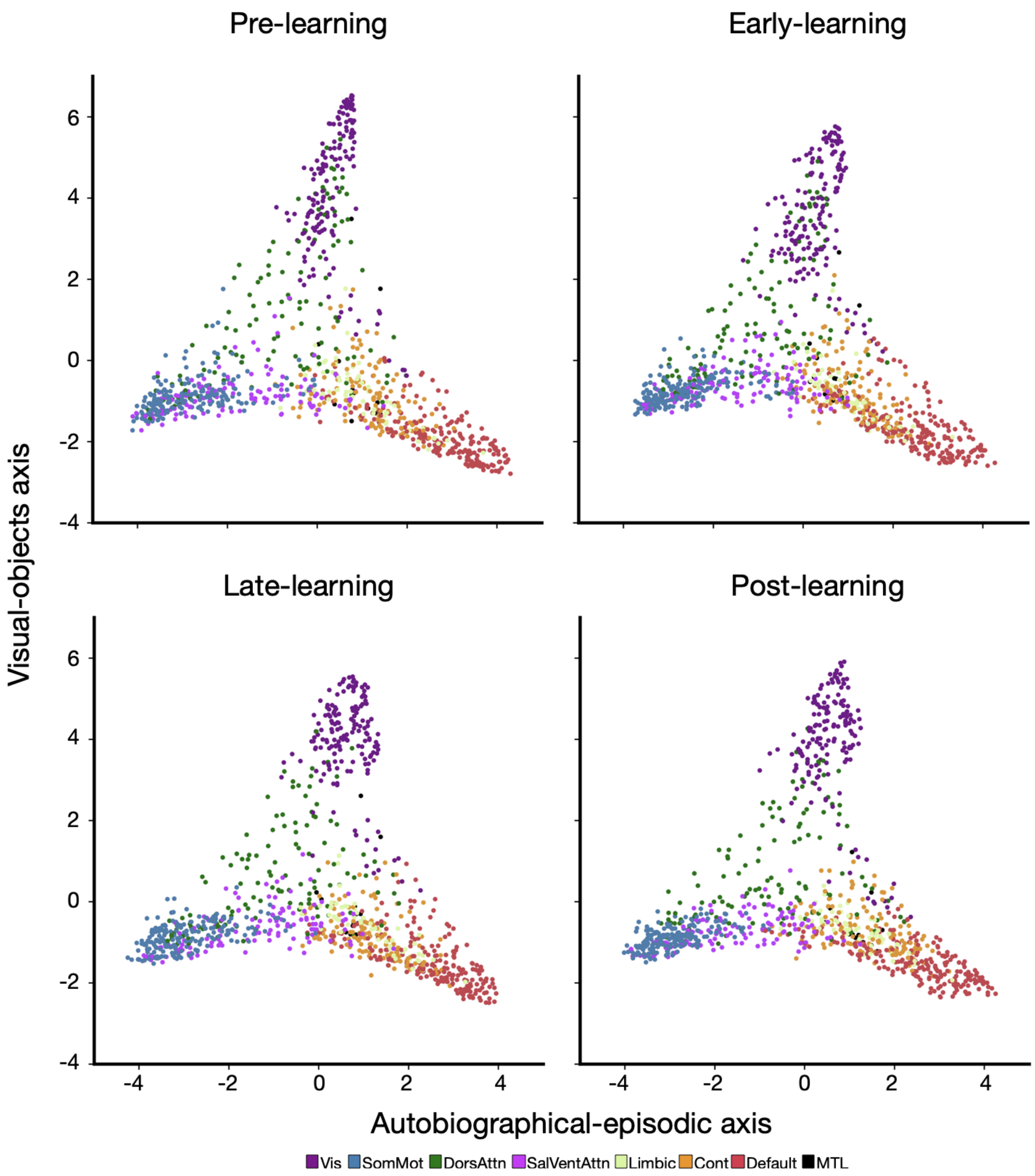


**Figure S3. Group average manifold connectivity across task epochs.** Functional organization of each task epoch of interest (Pre-, Early-, Late-, and Post-learning). Each brain region is depicted as a point in two-dimensional manifold space, and its colour determined by its functional network assignment, denoted at the bottom (30-31, 50).
